# Improved transcriptome assembly of *Monarda citriodora* reveals candidate HD-ZIP IV transcription factors involved in trichome development and secondary metabolism

**DOI:** 10.64898/2026.04.21.719861

**Authors:** Sonali Andotra, Kulsoom Shafeeq, Koushik Pal, Aasim Majeed, Prashant Misra

## Abstract

Members of the HD-ZIP transcription factor family play an important role in plant processes, including growth, development, metabolism, and stress response regulation. Among these, the sub-family IV members regulate epidermal cell differentiation, trichome development, and secondary metabolism. *Monarda citriodora*, an aromatic plant, produces economically important essential oils enriched in thymol. Thymol and other related monoterpenes are primarily biosynthesized and stored in glandular trichomes. Despite its significant economic value, the comprehensive identification of the transcription factor families has not been studied in this plant species. Given the importance of HD-ZIP-IV members in regulating trichome development and secondary metabolism, we identified these members in *M. citriodora* in the present study. To this end, firstly, we carried out transcriptome sequencing of *M. citriodora* flowers, and the resulting reads, along with previously sequenced reads, were used to reconstruct a transcriptome assembly. The assembled transcripts represented all major plant parts. Using this improved assembly, HD-ZIP-IV members were identified. Their expression profiles and phylogenetic positions, in conjunction with those of known regulators, identified candidate genes involved in the secondary metabolism and/or trichome development in *M. citriodora*. Furthermore, through gene co-expression analysis, several *McHD-ZIP-IV* members were found to be co-expressed with *McDXS* and *McTPS* genes. These *McHD-ZIP-IV* members may serve as key candidate genes for functional analysis to determine the regulation of trichome development in *M. citriodora*. Taken together, the present study provides a resource for improving *M. citriodora* using molecular tools.

## Introduction

The homeodomain-Zipper (HD-ZIP) family of transcription factors (TF), one of the plant-specific families, is characterized by a conserved homeodomain (HD). These transcription factors orchestrate diverse processes in plant biology, including the regulation of plant growth, development, metabolism, and stress adaptation. All members of this family contain an HD domain composed of 60 amino acids and an adjacent leucine LZ domain. The HD domain is responsible for DNA binding, and the LZ domain mediates protein-protein interactions for dimerization **(Xie et al, 2025; Ma et al, 2024).** Phylogenetically, HD-ZIP family members are classified into four sub-families (HD-ZIP I, HD-ZIPII, HD-ZIPIII, and HD-ZIP IV), which display distinct compositions of additional conserved domains. While the HD-ZIP I transcription factors possess a conserved HD domain and a less-conserved LZ domain, the members of the HD-ZIP II sub-family harbour an additional conserved CPSCE (Cys, Pro, Ser, Cys, and Glu) motif at their C-terminus. The domain composition in HD-ZIP III sub-family members is comparatively more elaborate due to the presence of additional conserved domains, namely the steroidogenic acute regulatory protein-related lipid transfer (START) domain, the START adjacent domain (SAD), and the MEKHLA domain. Thus, HD-ZIP III subfamily transcription factors display the HD-LZ-START-SAD-MEKHLA composition oriented from the N-terminus to the C-terminus. The HD-ZIP IV sub-family lacks MEKHLA domains but is otherwise similar to HD-ZIP III sub-family transcription factors in conserved domain composition **(Ariel et al. 2007; Zheng et al. 2025).**

HD-ZIP I and II TFs bind to similar pseudo-palindromic *cis*-acting elements, with consensus sequences being CAATNATTG. However, their central nucleotides differ with A/T and C/G for the binding of members of HD-ZIP I and HD-ZIP II subfamilies, respectively (Ariel et al, 2007). The HD-ZIP III TFs bind to the GTAAT(G/C)ATTAC sequence and HD-ZIP IV TFs bind to the TAAATG(C/T)A sequence, with a TAAA core sequence **(Henriksson et al, 2005).** Members of different subfamilies have been reported to regulate distinct processes. HD-ZIP I TFs play a role in the regulation of responses to stress, light, and hormones such as ABA, as well as in organ development, embryogenesis, and de-etiolation **(Gong et al., 2019).** The members of HD-ZIP II are mainly involved in auxin response, light signalling, and shade avoidance response **(Li et al, 2022).** The HD-ZIP III TFs have been implicated in the regulation of cell differentiation, vascular system development, shoot apical meristem, embryogenesis, lateral organ initiation, leaf polarity, etc. (**Elhiti & Stasolla, 2009; Ohashi-Ito et al, 2013).**

Members of the HD-ZIP IV subfamily primarily regulate epidermal cell differentiation and related biological processes in plants. To this end, the HD-ZIP-IV transcription factors play a crucial role in the initiation and development of trichomes in different plant species **(Schrick et al. 2023; Li et al. 2025).** Importantly, several secondary metabolites, such as specialized terpenoids and essential oils, are primarily biosynthesized and stored in glandular trichomes, whose regulation is controlled by a transcriptional network involving HD-ZIP IV subfamily transcription factors (**Schuurink & Tissier, 2019**; Kundan et al., 2020). For example, in *Artemisia annua*, HD-ZIP transcription factors HD1 and HD8 positively regulate the development of glandular trichomes. The glandular trichomes of *A. annua* are primary sites of the biosynthesis and storage of an antimalarial sesquiterpenoid, artemisinin. The overexpression of these TFs led to increased trichome density and enhanced artemisinin content in transgenic plants (**Yan et al. 2017; Yan et al. 2018).** In tomato, another HD-ZIP IV sub-family transcription factor, namely Woolly (Wo), regulates multicellular trichome development and differentiation **(Wu et al. 2023).** In *Arabidopsis thaliana*, the HD-ZIP IV transcription factor GL2 (GLABRA2) plays a crucial role in the positive regulation of unicellular trichome development. These transcription factors work in concert with other transcription factors to fine-tune trichome development. For example, in *A. thaliana*, GL2 functions in a complex with bHLH transcription factors and a WD40-repeat protein to regulate trichome development (Li et al. 2025). In *A. annua*, an R2R3 MYB transcription factor, MIXTA interacts with HD8 to regulate the expression of genes involved in cuticle and trichome development (Yan et al. 2018). A few HD-ZIP subfamily TFs directly regulate the biosynthesis of secondary metabolites such as anthocyanins and flavanols **(Wang et al. 2020).** In addition, the HD-ZIP IV transcription factors have also been shown to be involved in the biosynthesis of cuticular wax and cuticle **(Liu et al. 2025).** Given their crucial role in regulating trichome development, HD-ZIP-IV subfamily members are targets for genetic manipulation to improve the yield of trichome-specific metabolites.

*Monarda citriodora*, a member of the family Lamiaceae, is an economically important aromatic plant. The essential oil and extract from this plant species are sources of monoterpenes such as ץ-terpinene, carvacrol, thymol, and thymoquinone. This plant is used for flavouring various beverages, meat, and bakery products. The essential oil has been reported to exhibit anti-cancerous and antimicrobial activities **(Pathania et al. 2013; Di Vito et al. 2019).** The bioactivity of plant extracts and essential oils is attributed to their monoterpenoid constituents. The molecular basis of the biosynthesis of the monoterpenoid constituents of this plant species essential oil remains poorly understood. Earlier, our group, for the first time, developed a transcriptome assembly using RNA sequencing of the samples from the stem, root, and leaf. The assembly was used to identify candidate genes involved in the biosynthesis of the constituents of essential oils in *M. citriodora* **(Sharma et al. 2023).** In addition, our study also demonstrated that the glandular trichomes are the primary sites of the biosynthesis and storage of these compounds. One terpene synthase gene encoding ץ-terpinene synthase was functionally characterized and demonstrated to be involved in the biosynthesis of thymol **(Wajid et al. 2024).** The promoter region of the ץ-terpinene synthase is primarily active in the glandular trichomes, further confirming the trichome-specific biosynthesis of aroma molecules. 1-deoxy-D-xylulose-5-phosphate synthase (DXS) catalyse the entry step of the methylerythritol phosphate (MEP) pathway, which produces substrates to primarily support the biosynthesis of aromatic monoterpenoids. Four DXS genes have been identified and functionally characterized from *M. citriodora*. Among these, McDXS2 was shown to be involved in the biosynthesis of aromatic monoterpenoids in this plant species **(Sharma et al. 2025).** Further, the McDXS2 promoter region was found to be primarily active in the glandular trichomes. Despite this accumulating knowledge, the molecular mechanisms underlying factors associated with essential oil biosynthesis remain to be studied in greater detail. One major limitation to this end is the lack of a high-quality transcriptome and genomic resource for this plant species.

Against this backdrop, in the present study, we improved our already-developed transcriptomic resource by sequencing the transcriptome of *M. citriodora* flower tissue. To the best of our knowledge, no transcription factor family has been characterized from *M. citriodora*. Given the crucial role of the HD-ZIP IV family in regulating glandular trichome development and metabolism, the reconstructed assembly was used to identify these transcription factors. We identified the members of HD-ZIP TF family and classified them into 4 distinct sub-families based on its conserved domains and structural features. Additionally, phylogenetic and expression analyses were performed to narrow down potential candidates involved in secondary metabolism and trichome development. Furthermore, a co-expression analysis of these HD-ZIP TFs was performed with the TPS and DXS gene families to identify key HD-ZIP TFs co-expressed with these family members. Taken together, the present study develops a molecular resource of HD-ZIP IV family members, which will be useful for understanding the molecular regulation of terpenoid biosynthesis as well as trichome development in *M. citriodora*.

## Material and Methods

### Plant material

The *M. citriodora* plants were raised using the same strategy described in our previous studies **(Sharma et al, 2023, 2025; Wajid et al, 2024).** Briefly, the plants were grown under controlled conditions: 16 h of light and 8 h dark cycles and 22°CL±L1. Two-week old seedlings were shifted to pots containing an equal mixture of vermicompost, soil and soil-rite. Subsequently, these seedlings were grown in walking plant growth chamber (Percival Scientific, Perry, IA, USA) additional 11-weeks till flower maturation. Then the flower tissue was harvested in liquid nitrogen and stored in -80oC. RNA isolation and transcriptome sequencing

The RNA was isolated according to the methodology described by **Sharma et al, (2023).** The quality of the isolated RNA was checked by gel electrophoresis and quantified using Nanodrop spectrophotometer. Furthermore, the RNA integrity (RIN) was determined using the Agilent Bioanalyzer 2100. Approximate 3 micro grams of the high-quality RNA was used for cDNA libraries preparation using the Illumina RNA Sample Preparation Kit. Prior to paired-end sequencing, the quality of the library was evaluated using the Agilent 2100 Bioanalyzer and the Qubit 2.0 fluorometer. Sequencing was performed on Illumina Novaseq 6000 (Illumina Inc., CA, USA) with 150 bp insert size. The RNA for other plant tissues like different developmental stages of leaves, stem and root were isolated as per the method described by **Sharma et al. (2023).**

### Assembly and annotations

The quality of the paired-end raw reads corresponding to flower was evaluated using FastQC **(Andrews 2017).** FastP **(Chen, 2023**) was used to remove low-quality reads and adaptors. The high-quality reads from other tissues (leaf, stem, and root) were generated previously (Sharma et al. 2023). These reads were merged with high quality paired-end reads of flower to reconstruct a more inclusive and comprehensive de novo transcriptome assembly using Trinity assembler **(Grabherr et al. 2011).** The paired-end reads were aligned back to the de novo assembly using Bowtie2 **(Langmead and Salzberg 2012)** to estimate read-representation. Additionally, BUSCO **(Simao et al. 2015)** analysis was performed to check the quality of the assembly. Furthermore, the redundant sequences were removed using CD-HIT-EST **(Li and Godzik 2006**) with 0.95 sequence identity threshold. Finally, the non-redundant assembly was structurally annotated using the TRAPID pipeline **(Van Bell et al. 2013),** with Plaza 4.5 Dicot database as a reference. Functional annotations were performed by aligning the transcripts against multiple public databases including TAIR, Swiss-Prot and CDD, Gene Ontology (GO), KEGG, pfam, and InterPro databases.

### HD-ZIP gene family identification

We used the hidden Markov model (HMM) profiles of the homeobox domain (PF00046) and the leucine zipper (LZ) domain (PF02183) to evaluate the M. citriodora protein sequences using the HMMER search tool **(Potter et al, 2018).** The significant hits obtained from HMMER were further evaluated using InterProScan 5.68 **(Jones et al, 2014)** and SMART databases **(Letunic et al, 2021**) to validate the presence of HD and LZ domains. The molecular weights (MW), theoretical isoelectric points (pI), and average hydropathicity (GRAVY) values of the HDZs were analyzed using the online Expasy tool ProtParam (https://web.expasy.org/protparam). Subcellular locations were predicted using the online tool Plant-mPLoc **(Chou and Shen 2010).**

### Phylogenetic analysis

Multiple sequence alignment (MSA) of the identified McHD-ZIP proteins was performed using MUSCLE **(Edgar, 2004).** Then the Maximum Likelihood (ML) phylogenetic tree was constructed using IQTREE **(Minh et al, 2020).** The visualization and editing of the ML tree was done by iTOL (**Letunic et al, 2024).**

### Spatial expression analysis of the McHD-ZIP IV sub-family

The high quality paired-end raw reads from leaf, stem, root, and flower were aligned against the newly reconstructed reference transcriptome of M. citriodora using STAR **(Dobin et al, 2013).** The read count estimation per gene was performed using FeatureCounts **(Liao et al, 2014).** The TPM values of the predicted McHD-ZIP-IV members were used for comparing their expression patterns across different tissues. Furthermore, the in silico predicted expression was checked through qRT-PCR. A total of 15 McHD-ZIP-IV genes were for qRT-PCR validation. First the RNA was isolated from root (RT), stem (ST), flower (FL), and different developmental stages of leaves small leaf (SL), medium leaf (ML), large leaf (LL). Tissue specific cDNA libraries were constructed using the Neoscript 1st strand cDNA synthesis kit (Genes2Me, India). The Step One Plus™ Real-Time PCR system (Applied Biosystems, Waltham, MA, USA) was used for qRT-PCR. Each reaction consisted of a total volume of 10 µL with 5µL of 2L×LSYBR green ROX mastermix, 0.2 µL each of forward and reverse primer, 1 µL template and 3.6 µL of deionised water. The PCR program was set at 95°C for 10 min and 42 cycles of 95°C (15s) and 60°C (1 min), 72°C (7 min). The M. citriodora elongation factor (McELF) was used as a reference gene. The 2-ΔΔCt method **(Livak and Schmittgen 2001)** was used to estimate the relative fold change in expression. The sequences of the primers used in the qRT-PCR are given in supplementary file S1.

### Gene co-expression analysis of McHD-ZIP-IV, TPS, and DXS genes

A gene co-expression analysis was performed between McHD-ZIP-IV members, Terpene synthases (TPS), and 1-deoxy-D-xylulose-5-phosphate synthase (DXS) genes. First, the hmm models of TPS N-terminal domain (PF01397) and Terpene synthase family, metal binding domain (PF03936) were used to identify the TPS proteins from *M. citriodora* proteome using HMMER scan. An e-value threshold of 1e-3 was used. The resulting hits were verified from SMART database. The four *McDXS* (McDXS1-4) identified from our previous study **(Sharma et al, 2025)** were also used for the co-expression analysis. The corresponding homologs of these *McDXS* members in the reconstructed assembly were first identified using blastx and then used for co-expression analysis. The co-expression network was then constructed using BIONERO (**Almeida-Silva & Venancio, et al, 2022).**

## Results

### Development and annotation of an improved transcriptome assembly of *M. citriodora*

*De novo* assembly of high quality paired-end reads by Trinity yielded a total of 377639 transcripts. Filtering of transcripts <300 nt length resulted into a total of 280399 transcripts. Removal of sequence redundancy by CD-HIT-EST generated a high quality non-redundant assembly consisting of 220881 transcripts (Table 1). Compared to our previous assembly with 191638 transcripts constructed from paired-end reads of leaf, stem, and root only, the addition of reads from flower has significantly improved the de novo assembly. All to all blast of these two assemblies revealed that a total of 189972 unique previous transcripts have significant hits against the new assembly, with 1666 missing transcripts. A total of 123682 newly assembled transcripts had at least one significant hit with the previous assembly, with 97199 transcripts additionally captured when flower tissue was incorporated. BUSCO assembly completeness analysis revealed that out of the 1614 queried genes, 1582 genes were retrieved successfully indicating 98.01% completeness.

**Table 1:**
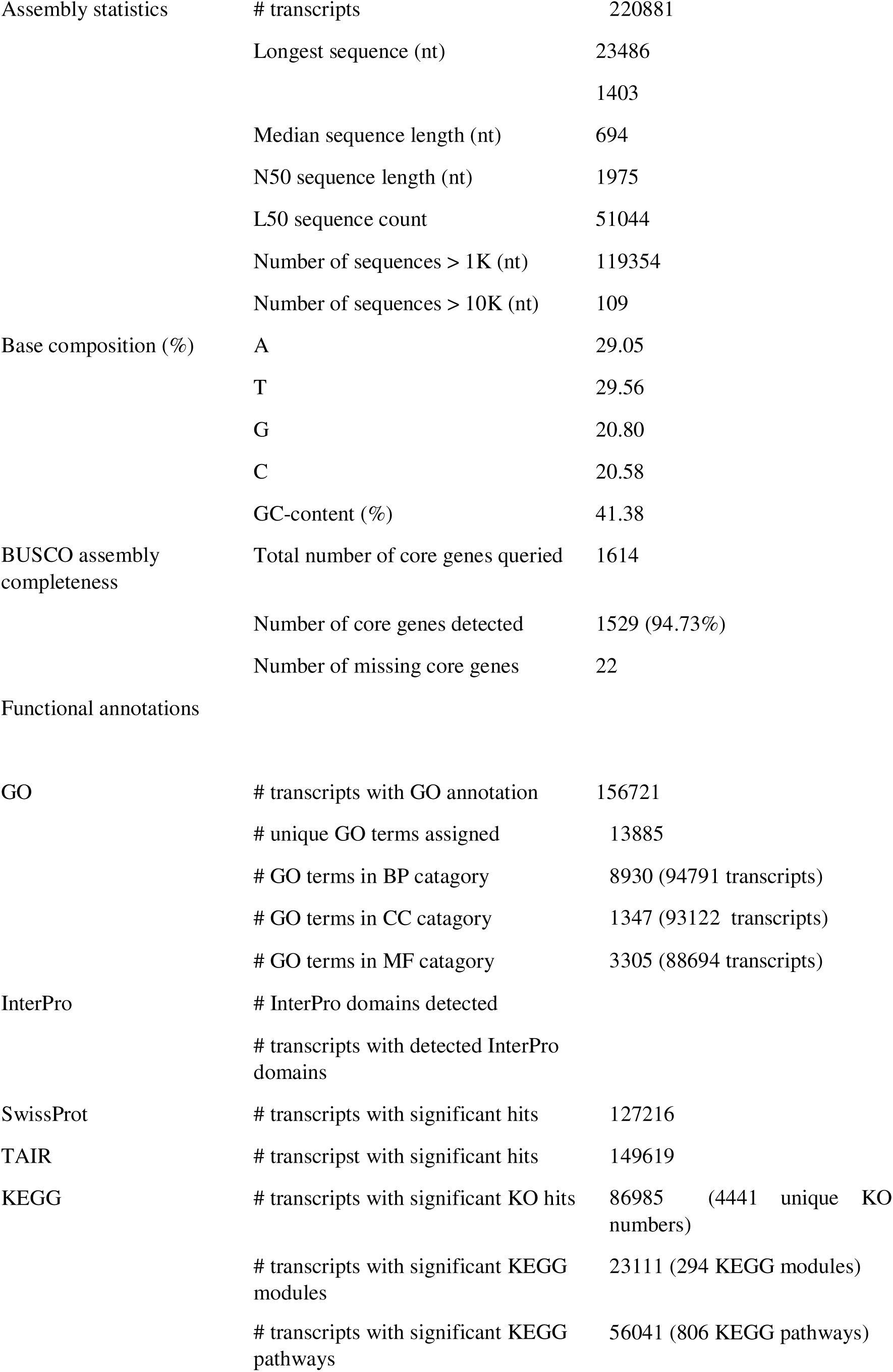
Summary statistic of assembled transcripts and annotations.

The functional annotation of the transcripts was performed by homology searches against different databases. A total of 149619 unique transcripts were found to have significant hits with the 23226 unique proteins in the TAIR database. 127216 unique transcripts have significant hits with 17067 unique proteins inthe SwissProt database, unique transcripts with the InterPro database, and unique transcripts with the pfam database. Furthermore, using InterProScan v5 (Jones et al, 2014), 137892 unique transcripts were found to have hits in the PANTHER database, 30951 in the CDD database, 111865 in the pfam database, and 135244 in the InterPro database (Figure 1). Moreover, GO annotation using eggNog revealed 156721 transcripts with 13885 unique GO terms. A total of 94791 transcripts have 8930 GO terms associated with biological processes (BPs), 93122 transcripts with 1347 GO terms associated with cellular components (CC), and 88694 transcripts with 3305 GO terms associated with molecular functions (MFs). Similarly, KEGG annotation revealed 86985 transcripts with 4441 unique KO terms, 23111 transcripts with 294 KEGG module terms, and 56041 transcripts with 806 unique KEGG pathway terms. The top 10 GO BPs, CC, and MFs and top 10 KEGG Kos, Modules, and Pathways are shown in figure 2.

**Figure 1:**
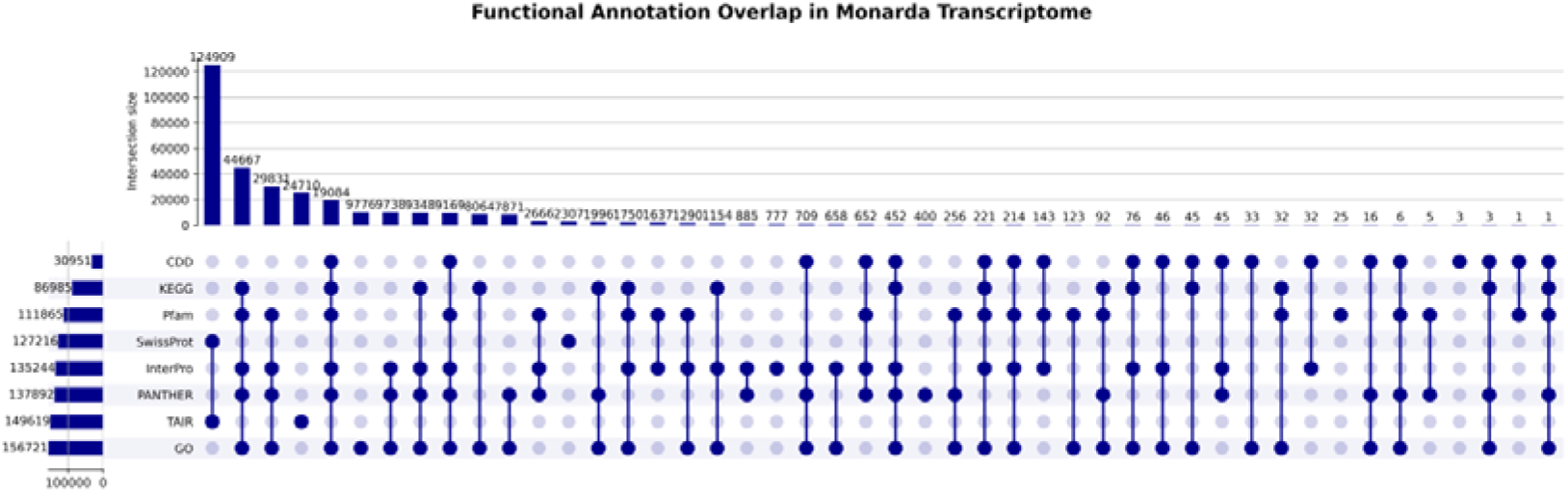
Summary of functional annotation of *M. citriodora* reference transcriptome assembly

**Figure 2:**
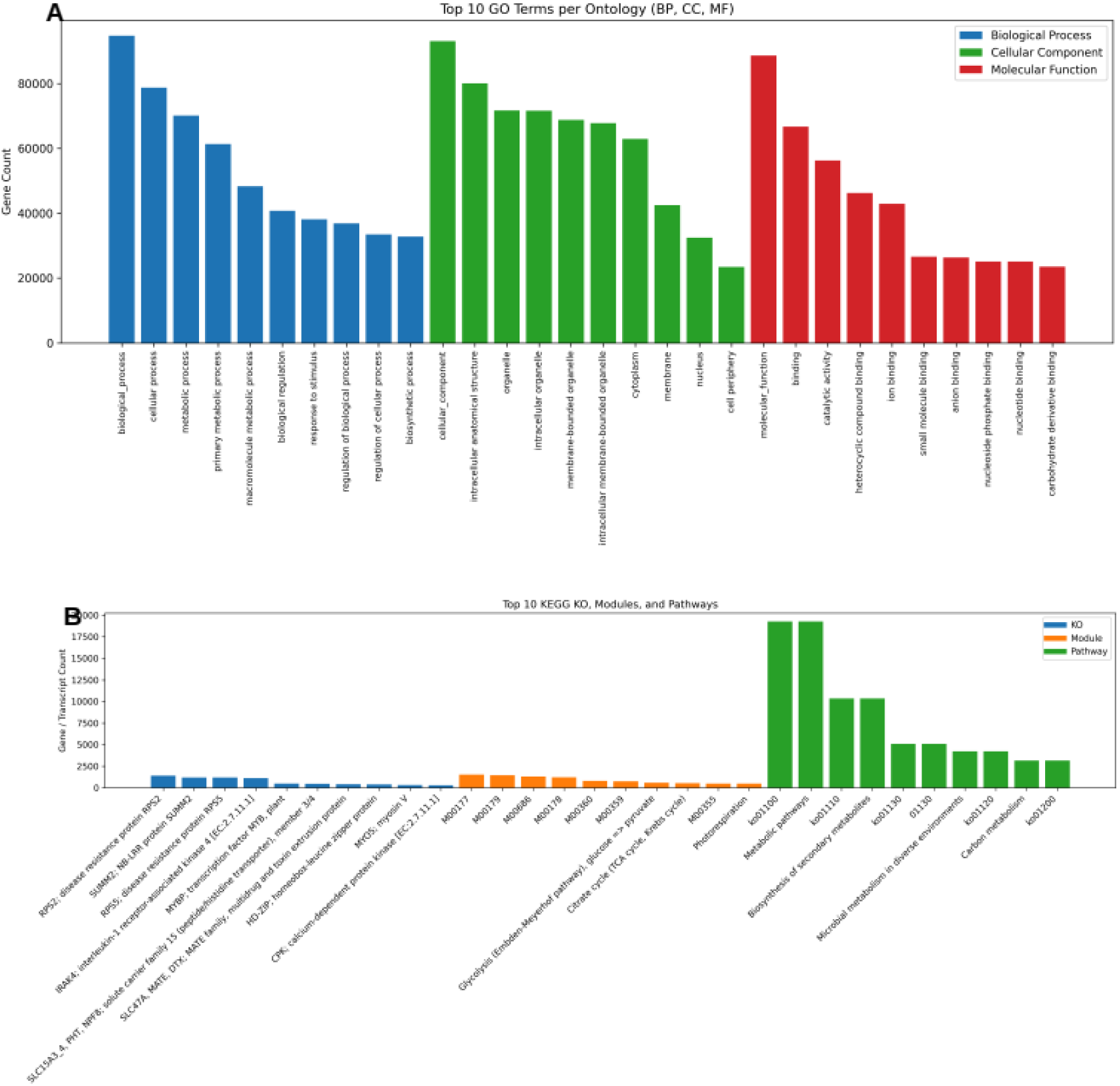
Top enriched GO (A) and KEGG (B) pathways

### HD-ZIP gene family identification, classification, and phylogeny

A total of 148 McHDZ-ZIP proteins were identified using HMMER search followed by SMART database verification. These proteins were classified into 4 groups (McHD-ZIP-I to McHD-ZIP-IV). The results revealed 97 proteins classified as McHD-ZIP-I, 8 as McHD-ZIP-II, 2 as McHD-ZIP-III, and 41 as McHD-ZIP-IV. Among the McHD-ZIP-IV subfamily members, 19 are either 5’ partial or 3’ partial and 22 were full length proteins (Table 2). Given the standard protein length range of 600 to 850 aa for HD-ZIP-IV members, we found a total of 10 McHD-ZIP-IV members that pass this-criteria. For maximum-likelihood (ML) based phylogenetic analysis were used additional 100 protein sequences from different species including functionally characterized HD-ZIPs in *Artemesia annua, Zea mays*, *Gossipyum hirsutum, Solanum lycopersicum*, as well as genome-wide HD-ZIPs of Oryza sativa and *Arabidopsis thaliana*. The AtMYB15 was used as an outgroup. A total of 3035 alignment length was generated following MSA by Muscle, with 1742 distinct patterns, 1143 parsimony-informative, 626 singleton sites, and 1266 constant sites. The ModelFinder implemented in IQTREE tested up to 1232 protein models, which revealed VT+F+R6 as the best model fitting our data based on BIC. ML phylogenetic tree revealed 4 major clades corresponding to HD-ZIP groups (Figure 3).

**Table 2:**
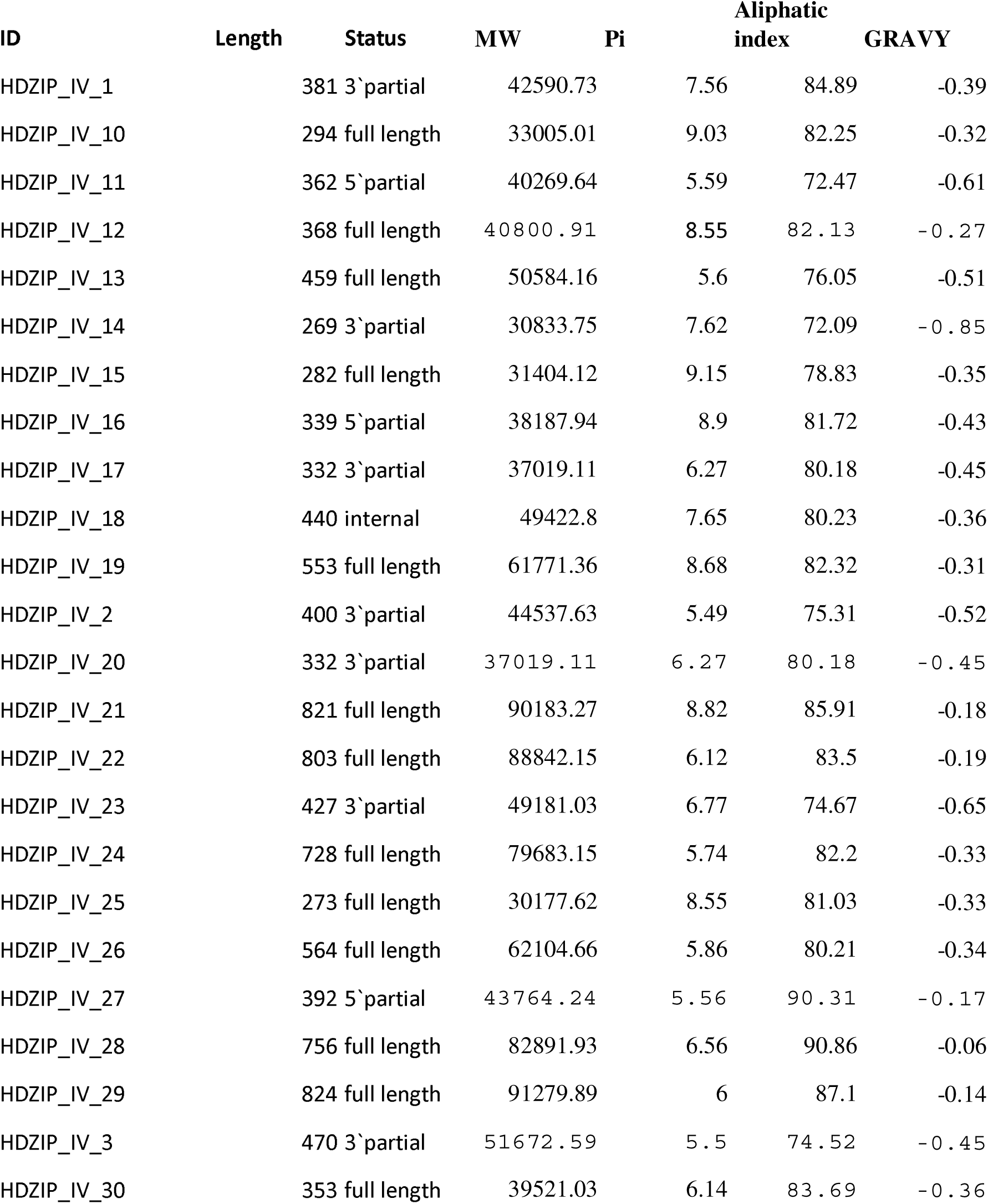
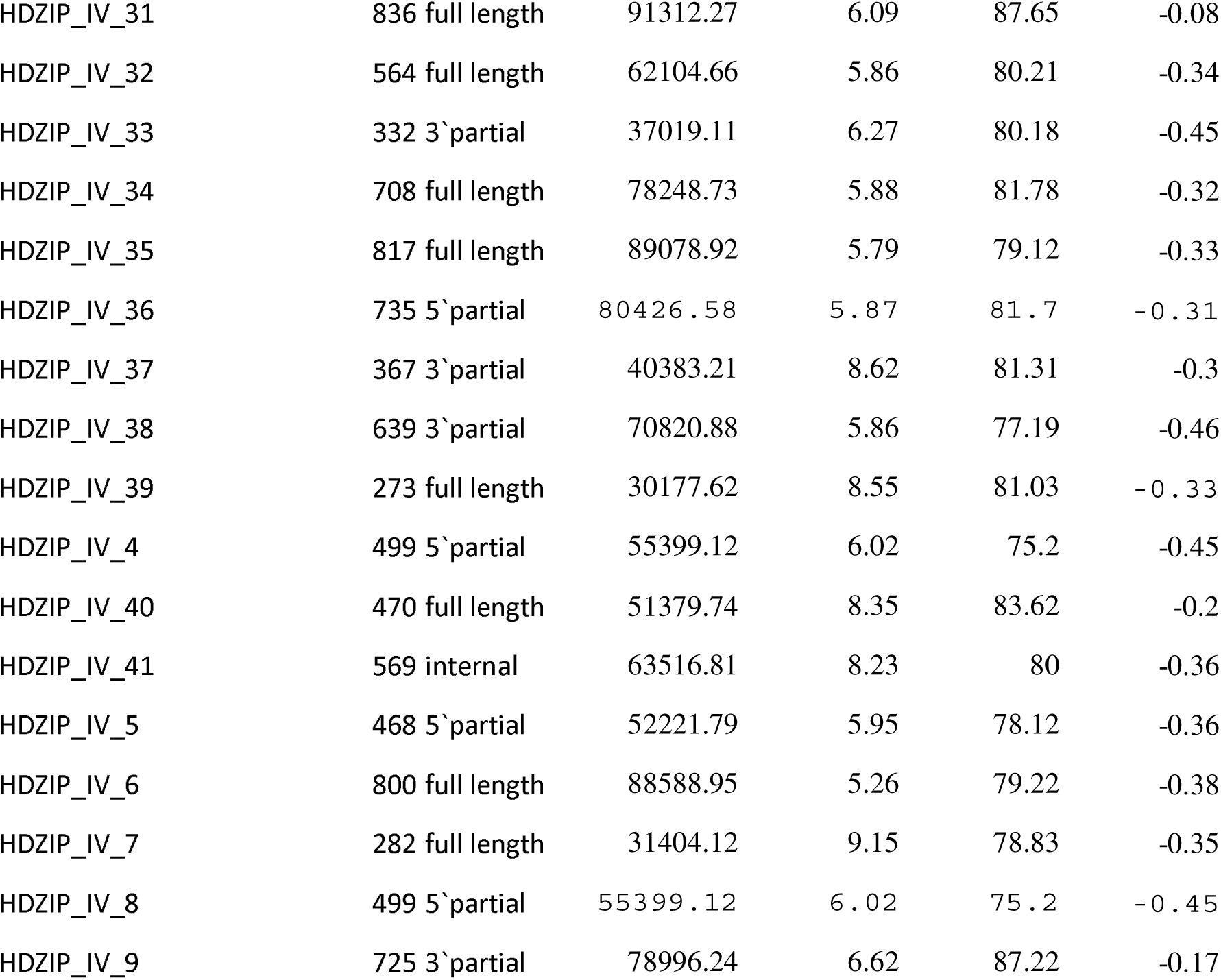
Physiochemical properties of the HD-ZIP-IV family TFs.

**Figure 3:**
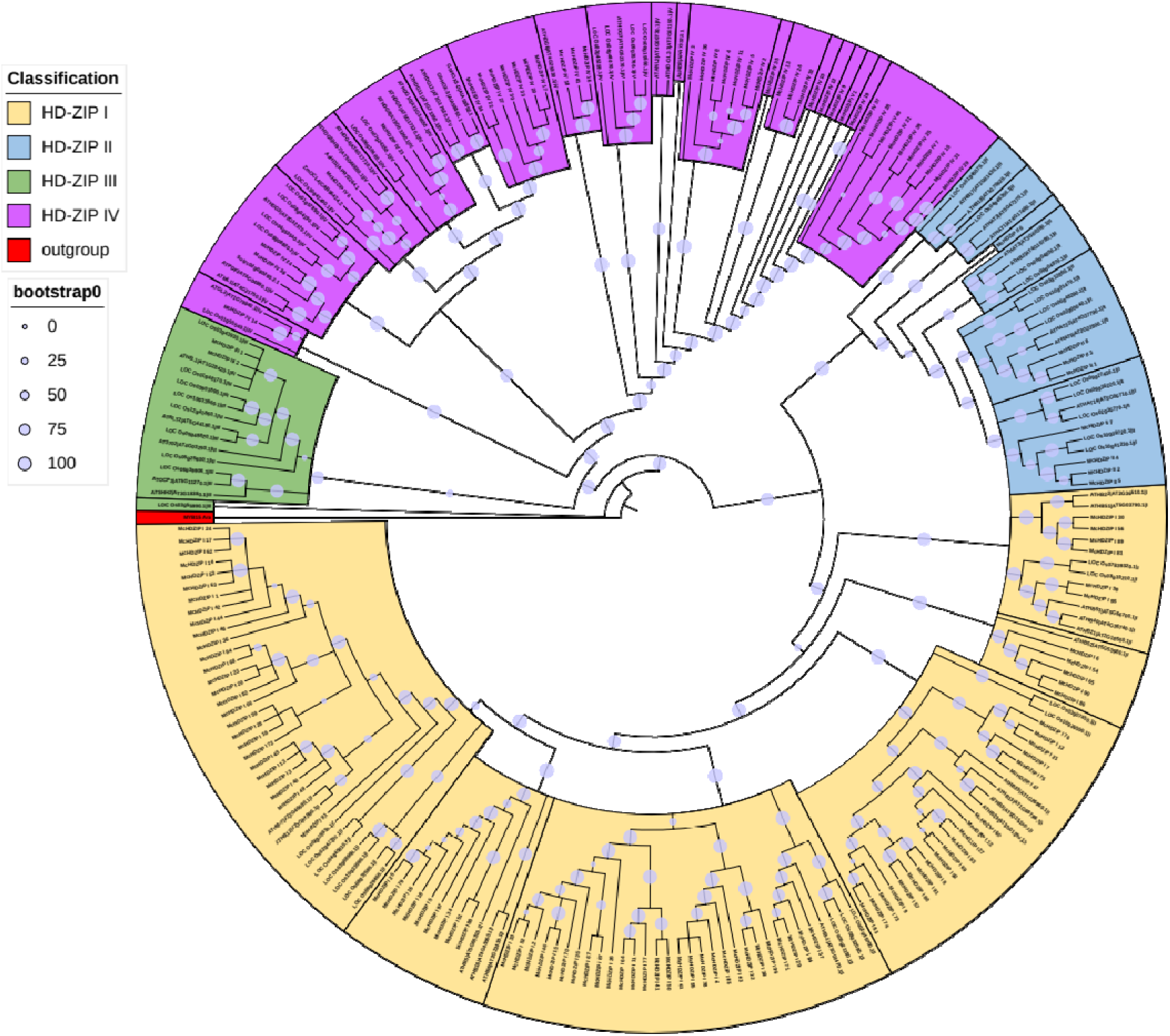
ML phylogenetic tree of McHD-ZIP TF family.

### Protein motif analysis of HD-ZIP-IV members

The protein sequences of the McHD-ZIP-IV proteins were used for prediction of conserved motifs using MEME suite v5.5.9 (accessed 10-April-2026) (Bailey and Elkan, 1994) (Supplementary file S2). Conserved motif analysis using MEME identified 20 motifs among the 41 HD-ZIP IV proteins. Motif-2 and motif-6 were present in all sequences and these motifs correspond to the core HD and START domains. Motif-1 (HD) and motif-4 (START) were detected in more than 95% of the proteins, whereas the remaining motifs were present only in subsets of sequences, indicating potential functional diversification.

### Expression profiling of *M. citriodora* HD-ZIP IV family

Spatial expression analysis of *M. citriodora* HD-ZIP IV family revealed 6 unigenes with highest expression in stem and root each, 4 genes with highest expression in flower, 2 with highest expression in L1, 3 with highest expression in L2, and 5 with highest expression in L3 (Figure 4). Furthermore, there were around 11 genes with overall higher expression across different leaf stages compared to other tissues. The functionally characterized AaHD1 was found to be involved in regulation of trichome development. The closest ortholog of *M. citriodora* corresponding to AaHD1 was found to be McHD-ZIP IV 6, whose gene expression was found to be highest in flower. Similarly, the closest orthologs corresponding to this AaHD8 in *M. citriodora* were McHD-ZIP IV 3, McHD-ZIP IV 36, McHD-ZIP IV 8, McHD-ZIP IV 4, McHDZIP IV 11, McHD-ZIP IV 5, and McHD-ZIP IV 2. The expression profiling of these genes revealed their relatively higher expression in different leaf stages compared to other tissues. These results agree well with the qRT-PCR results (Figure 5).

**Figure 4:**
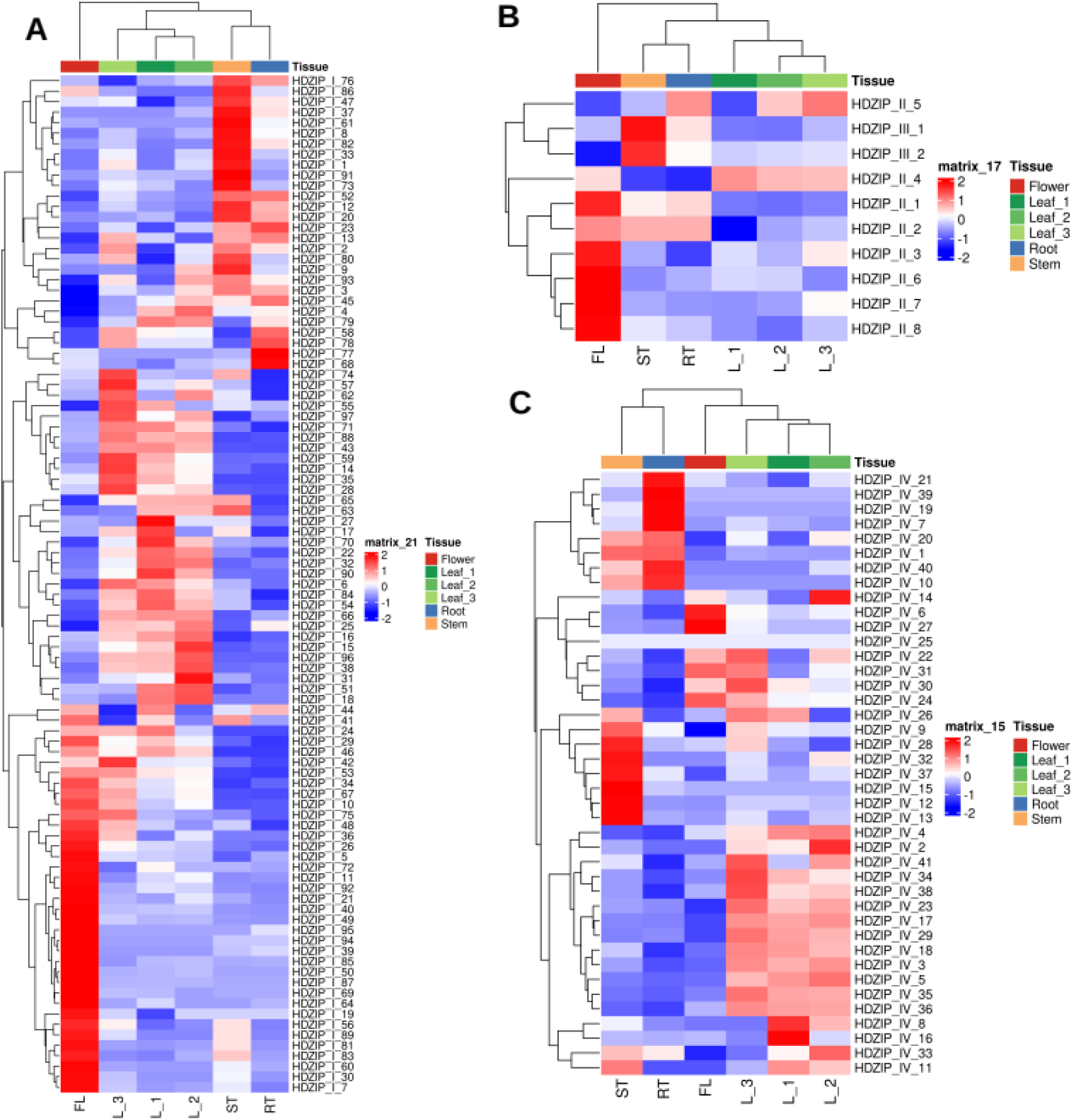
Heat maps showing expression pattern across different tissues of *M. citriodora.* (A) Expression profile of McHD-ZIP-I family. (B) Expression profile of McHD-ZIP-II & III familIes. (C) Expression profile of McHD-ZIP-IV family.

**Figure 5:**
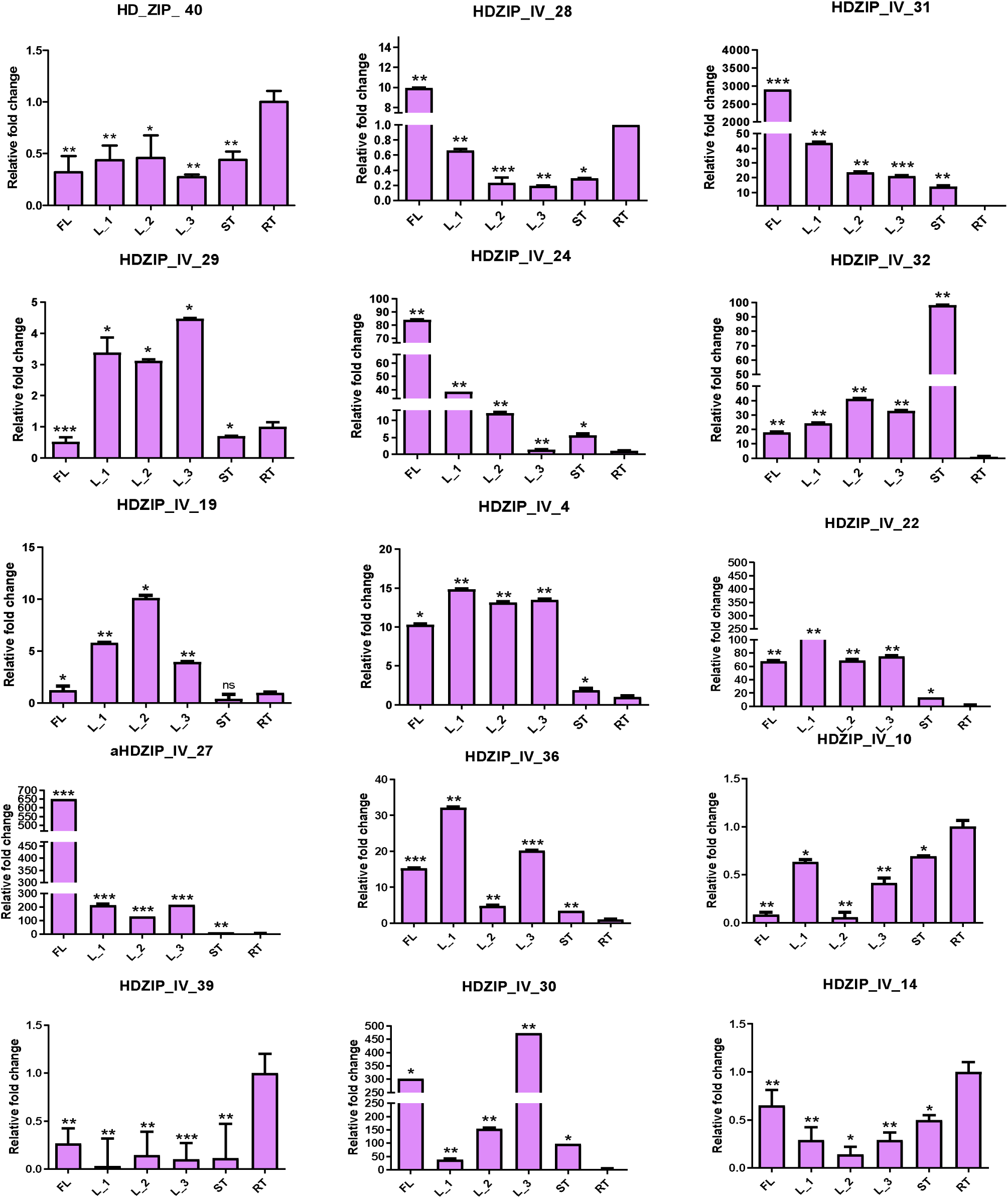
qRT-PCR based expression analysis of some selected *HD-ZIP* IV genes. The relative expression values are expressed in the form of relative fold-change with respect to the expression in the root tissue. FL, flower, L1, L2 and L3, different developmetal stages of leaves, ST, stem and RT, root

### Gene co-expression analysis of *McHD-ZIP-IV*, *TPS*, and DXS genes

DXS and TPS genes play crucial role in the biosynthesis of diverse specialized terpenoids including monoterpenes. Therefore, in order to identify the potential connection between HD-ZIP IV genes, DXS and TPS genes, a co-expression analysis was carried out. To this end, firstly, we identified complete repertoire of TPS gene family using the reconstructed transcriptome resource.

First, identification of the members of TPS and DXS gene family members was done to perform their gene co-expression analysis with the McHD-ZIP-IV. The HMMER scan against *M. citriodora* protein using the TPS N-terminal domain (PF01397) and Terpene synthase family, metal binding domain (PF03936) resulted into 83 TPS family members (McTPS_01 – McTPS_83). In order to phylogenetically classify these TPS proteins, 41 additional TPS proteins of *Arabidopsis thaliana* and 47 functionally characterized TPS proteins in different species were used (Wajid et al, 2024). The results revealed that McTPS genes clustered into 7 different clades (clade a-g) (Figure 6). The TPS-a includes sesqueterpene synthases and our analysis showed 23 McTPS members clustered within this clade. A total of 2 McTPS clustered into clade-b, suggesting that these TPSs could encode important monoterpene or isoprene synthases (Wajid et al, 2024). The TPS-c subfamily, which is most conserved among land plants, include important members such as copalyl diphosphate synthase, ent-kaurene synthase, and other diterpene synthases that are einvolved in crucial steps of the gibberellin biosynthesis, which regulates developmental and processes like stem elongation, seed germination, and flowering time (Li et al, 2025). A total of 10 McTPS members clustered within TPS-c clade. The TPS-d is gymnosperm specific and our analysis showed no McTPS clustering within this clade. The results of this study also showed that 10 and 5 McTPS members clustered into TPS-e and TPS-f, respectively. Both TPS-e and TPS-f include kaurene synthase and other diterpene synthases, monoterpene, and sesqueterpene synthases. A total of 14 McTPS members clustered with TPS-g that includes members from sesqueterpene, monoterpene and diterpenes synthases (Chen et al, 2011).

**Figure 6:**
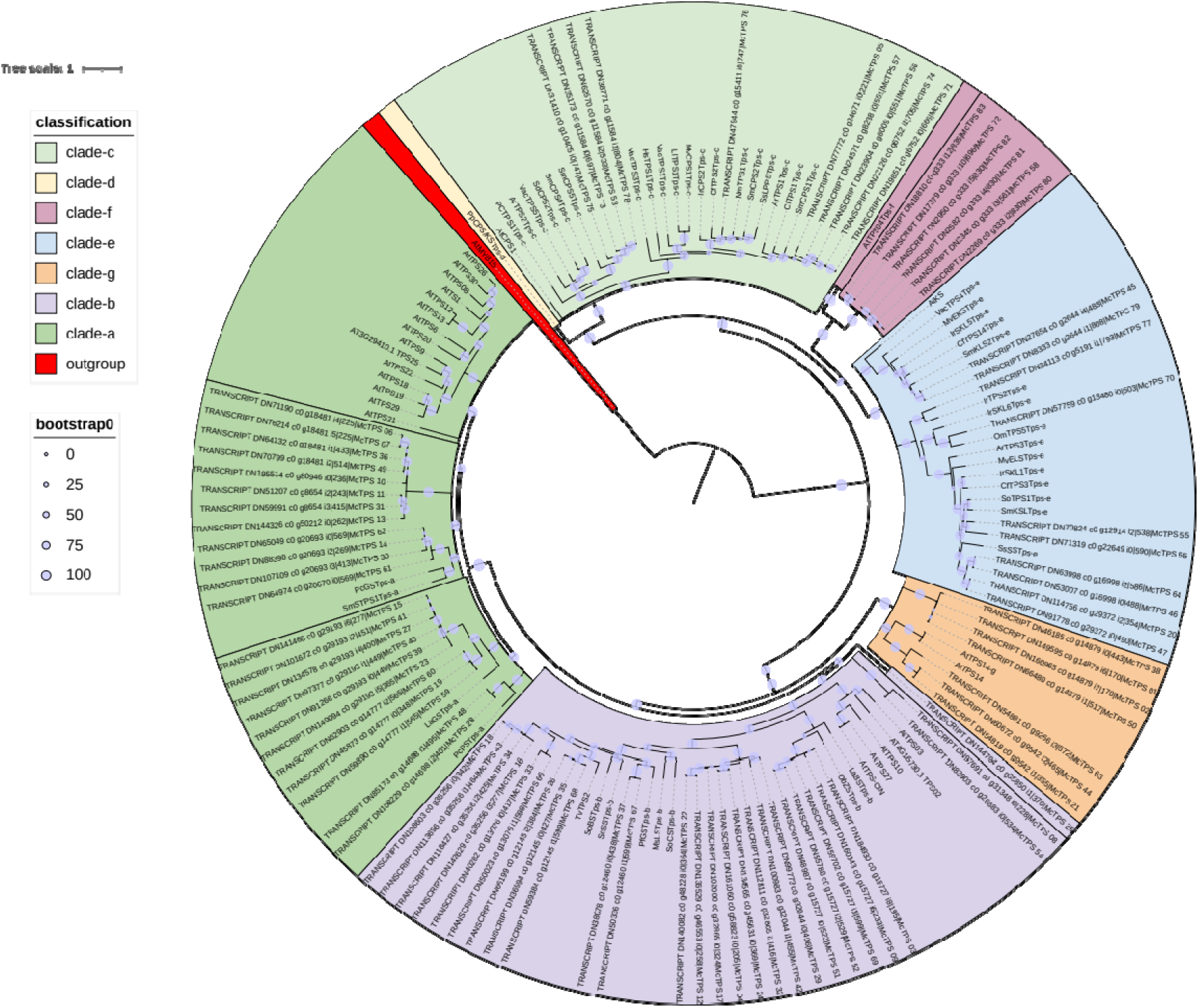
ML phylogenetic tree of *McTPS* gene family.

Next, the TPM based expression matrix of McHD-ZIP-IV, McTPS, and the McDXS members, constituting a total of 144 unigenes, was used as an input for gene-coexpression analysis. A total of 3 modules were detected using BIONERO, among which the modules turquoise and brown were highly positively (R^2^ = 0.75, p-value < 0.01) and negatively (R^2^ = - 0.81, p-value < 0.001) correlated with the flower, respectively. Similarly, the module blue was strongly correlated with the root (R^2^ = 0.76, p-value < 0.01) (Figure 7). Given the correlation threshold of R^2^ >= 0.8 & p-value <= 0.05, we observed that the blue module genes McHD-ZIP_IV_39 and McHD-ZIP_IV_19 were strongly correlated with McTPS_17 and McTPS_24 (Figure 8). These McTPS genes belong to the TPS-b clade, which are especially involved in the monoterpene biosynthesis. Since the blue module showed significant positive correlation with root, it is possible that these McTPS and McHD-ZIP genes are involved in the monoterpene metabolism in the roots of *M. citriodora*. In the brown module, McHD-ZIP_IV_4 was found to be negatively correlated with the McTPS_33. This module shows strong negative correlation with the flower,it may be possible that the monoterpenoid metabolism in the flower is negatively regulated by this McHD-ZIP by controlling the expression of McTPS_33. Similarly, in the torquoise module, McHD-ZIP_IV_27 and McHD-ZIP_IV_06 exhibited strong positive correlation with McTPS_08, McTPS_09, McTPS_25, McTPS_35, McTPS_52, McTPS_67, and McTPS_69. Except the McTPS_08, all of these McTPs genes along with McTPS_51 were also positively correlated with McHD-ZIP_IV_24. All these McTPS belong to TPS-b clade. Additionally, except the McTPS_08, all these McTPS genes also showed significant positive correlation with McDXS1 and except McTPS_25, all exhibited significant positive correlation with McDXS4. Moreover, McDXS3 was positively correlated with McTPS_67. Since the turquoise module exhibited strong positive correlation with the flower and the said McTPS belong to the TPS-b clade, it may be possible that monoterpene metabolism in the flowers of *M. citriodora* may be controlled by these McHD-ZIP by controlling the expression these corresponding McTPS and McDXS genes.

**Figure 7:**
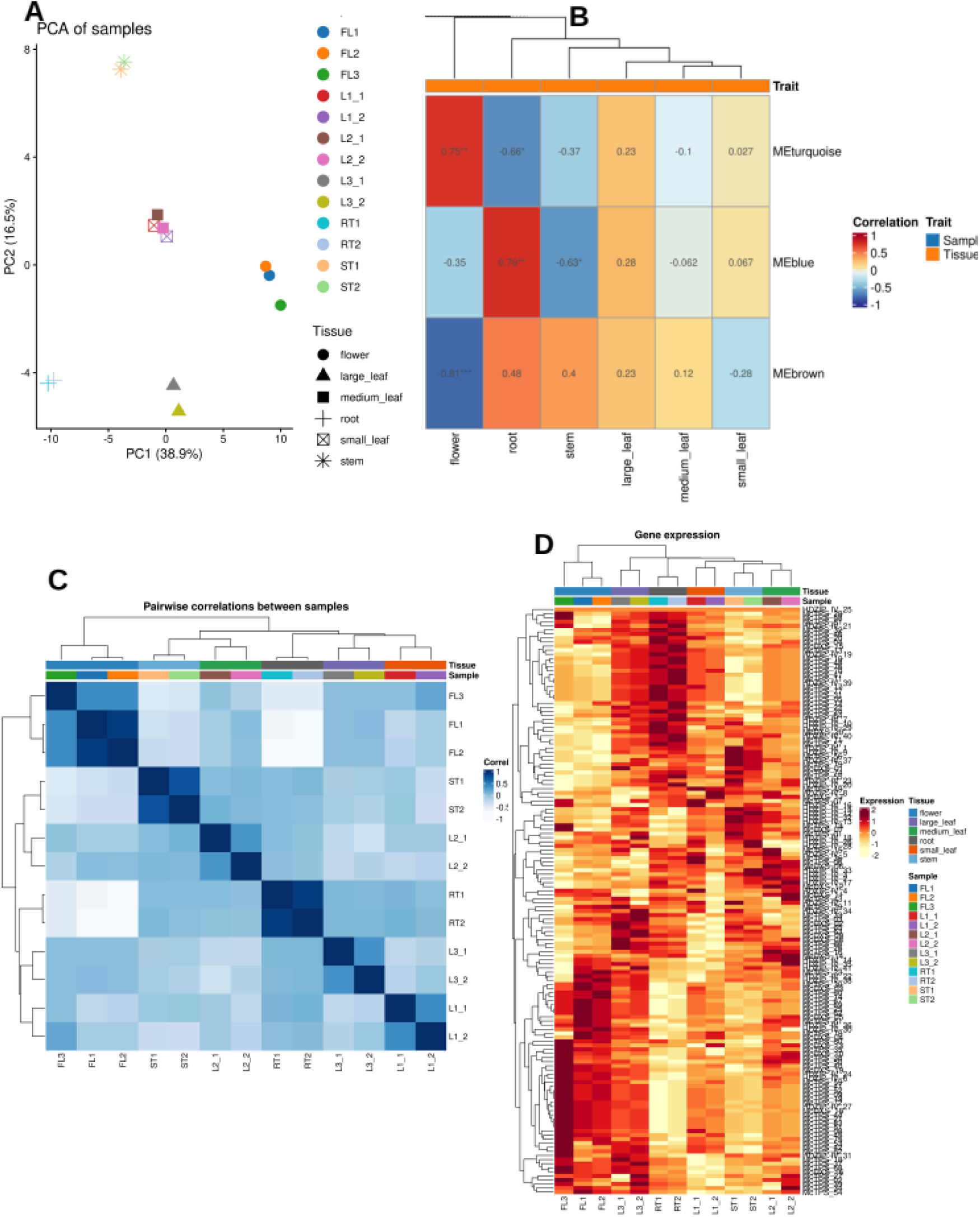
Summary of gene co-expression analysis. (A) Principal component analysis chart. (B) Heatmap showing tissue-trait relation. (C) Sample correlation heatmap. (D) Heatmap of genes used for gene-coexpression analysis.

**Figure 8:**
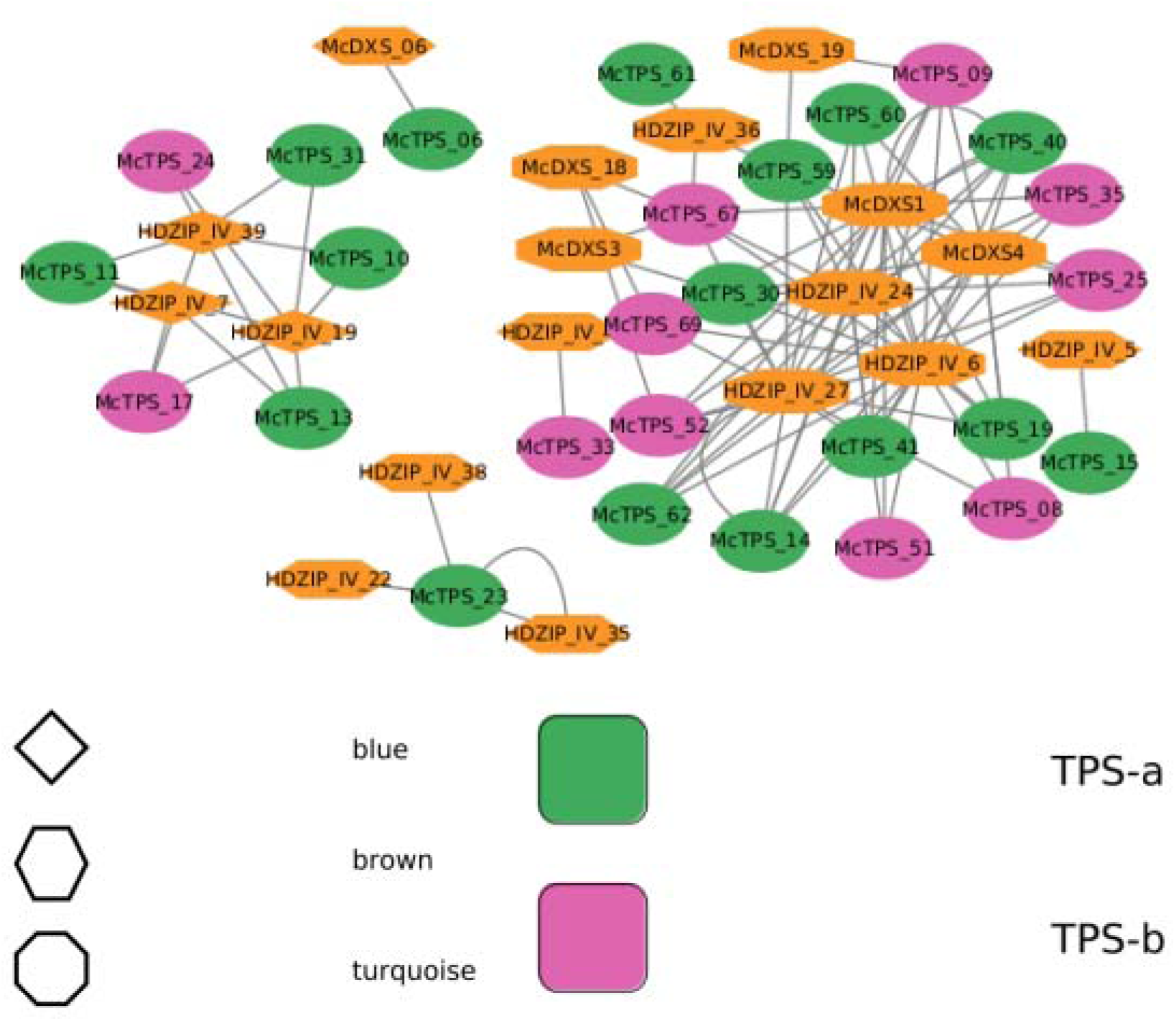
Co-expression network based on significant correlations among McHD-ZIP, *McTPS*, and *McDXS* members. The green circles represent McTPS-a clade while the pink circles represent McTPS-b clade. Members in the blue, brown, and turquoise modules are represented as diamond, hexagon, and octagon shapes, respectively.

Furthermore, among the 9 McTPS genes belonging to blue module and clustering with the TPS-a clade, the McTPS10, McTPS11, McTPS13, and McTPS31 exhibited strong positive correlation with McHD-ZIP_IV_39 and McHD-ZIP_IV_19. Since these genes correspond to the blue module that showed positive correlation with the root and TPS-a clade, which is mostly involved in the sesqueterpene metabolism, these McHD-ZIPs may govern the latter in the roots of *M. citriodora* by regulating the expression of these McTPS genes. Similarly, among the 5 McTPS genes of the TPS-a clade and belonging to the brown module, it was observed that McTPS_15 was positively correlated with McHD-ZIP_IV_5 & that of McTPS_23 was positively correlated with McHD-ZIP_IV_38, McHD-ZIP_IV_22, and McHD-ZIP_IV_35. Additionally, the McDXS1 was significantly positively correlated with HDZIP_IV_1, HDZIP_IV_10, HDZIP_IV_20, HDZIP_IV_29, and HDZIP_IV_40. Since the brown module was negatively correlated with the flower, it appears that thes McTPS, McDXS, and McHD-ZIPs operate outside the flower to regulate sesqueterpene metabolism. Likewise, a total of 9 McTPS genes belonging to turquoise module clustered with the TPS-a clade. This module was highly correlated with the flower and interestingly, 8 of those 9 genes showed strong positive correlation with McHD-ZIP_IV_27, McHD-ZIP_IV_24 and McHD-ZIP_IV_24, while the McHD-ZIP_IV_6 showed high positive correlation with 7 of those McTPS genes. Additionally, McHD-ZIP_IV_24, McHD-ZIP_IV_6, and McHD-ZIP_IV_27 also exhibited significant positive correlation with McDXS4 and McDXS1. Since the turquoise module exhibited a significant positive correlation with the flower, these results suggest that the said McHD-ZIPs, McDXS, and McTPS-a genes may be involved in the sesqueterpene metabolism in the flower of *M. citriodora*.

## Discussion

*M. citriodora* produces economically important essential oil enriched in monoterpenes. Despite its significant economic importance, the molecular studies focussing on the identification of transcription factors have not been carried out so far. Given the importance of HD-ZIP IV transcription factor in the regulation of growth, development and metabolism, especially in the development of glandular trichome, in the present study, we carried out transcriptome-wide analysis and characterization of HD-ZIP IV family members in *M. citriodora*.

We had previously generated transcriptome assembly of *M. citriodora* using paired-end reads from three tissues viz leaf, stem, and root, which yielded a total of 191638 non-redundant transcripts with transcript length cut-off of ≥ 300 nt **(Sharma et al, 2023).** However, this assembly lacked representation of transcripts from flowers. For the comprehensive analysis of gene families aimed at the functional genomics, it is imperative to develop a basic transcriptome assembly, which represent all major plant parts. Therefore, in this study, the paired-end RNAseq reads from the *M. citriodora* flowers were developed. These reads were assembled with the previously sequenced reads from leaf, stem, and root to reconstruct a more comprehensive *de novo* transcriptome assembly. Compared to our previous assembly, the addition of reads from the flowers has significantly improved the *de novo* assembly. All to all blast of these two assemblies revealed that a total of a total of 123682 transcripts had at least one significant hit with the previous assembly, with 97199 transcripts additionally captured when flower tissue was incorporated. The BUSCO completeness analysis also showed improvement over previous assembly with 98.02% completeness compared to 97.5% with the previous one (Sharma et al. 2023). Furthermore, the N50 was also improved from the previous 999 to current 1975. Additionally, the read-representation analysis also showed improved read alignment of 93.06% with the new assembly compared to 87% with the previous assembly. These parameters show significant improvement over the previous assembly. The new assembly, thus, represents a more comprehensive reference transcriptome resource for the functional genomics of *M. citriodoara*.

Transcriptome-wide analysis of the HD-ZIP family led to the identification of a total of 148 McHD-ZIP members in *M. citriodora*. As compared to the number of HD-ZIP genes, in other plant species with diploid genome, for example *Arabidopsis* (48 genes), and *Helianthus annuus* (55 genes), *Hordeum vulgare* (32 genes), and *Oryza sativa* (33 genes) the number of identified genes McHD-ZIP genes is significantly higher in *M. citriodora* **(Agalou et al. 2008; Ahangarani Farahani et al. 2025; Li et al. 2019).** Since HD-ZIP sub-family IV plays the major role in the development of glandular trichomes and thereby influence the yield of secondary metabolites, we focussed our study on the HD-ZIP subfamily IV members. The HDZIP-IV family members have been identified in different plant species. For example, the genome wide analysis led to the identification of 16, 17, 11, 21 HD-ZIP-IV genes in Arabidospis, maize, rice and banana, respectively (**Nakamura et al. 2006; Javelle et al. 2011; Pandey et al. 2016).** In the present study, comparatively higher number of HD-ZIP IV members has been identified in *M. citriodora*. This expansion in the number of genes may be attributed to the gene family evolution involving whole genome duplication. Further, the increased number may be due to the presence of truncated transcripts originating due to the short-reads. Further refinement of the assembly using third next-generation sequencing may resolve this issue. Nevertheless, the present work develops a database of HD-ZIP-IV genes for functional analysis.

HD-ZIP-IV family members are primarily involved in the regulation of epidermal cell differentiation in plants. Of special interest are the epidermal appendages, glandular trichomes, which are the primary sites of the biosynthesis and accumulation of secondary metabolites. Several HD-ZIP-IV members have been identified as positive regulators of glandular trichome development in diverse plant species such as tomato, tobacco and *A.annua* (**Schrick et al. 2023; Li et al. 2025).** Certain members of the family Lamiaceae such as Mentha, *Thymus vulgaris*, *Monarda citriodora* are rich source of economically important essential oils. Like other secondary metabolites, the essential oils are primarily produced in glandular trichomes **(Qamar et al. 2022; Sharma et al. 2023).** However, molecular regulators of the development of the trichomes in the family Lamiaceae remains poorly studied. Nevertheless, based on the studies on glandular trichome development in plant species tomato, tobacco and *A. annua*, it can be speculated that regulators of glandular trichome development are conserved across diverse plant species. To this end, one HD-ZIV IV member has been shown to positively regulate glandular trichome development in *Pogostemon cablin*, an essential oil producing plant from the family Lamiaceae **(Xie et al. 2025).** In *A. annua*, two HD-ZIP-IV family transcription factors, namely HD1 and HD8 play pivotal role in the glandular trichome development. The genetic manipulation of these genes affects the content of artemisinin in *A. annua* **(Yan et al. 2017; Yan et al. 2018).** Our phylogentetic analysis led to the identification of putative orthologs of HD1 and HD8 in *M. citriodora*. The phylogenetic analysis revealed one *M. citriodora* HD-ZIP-IV gene (McHD-ZIP-IV-6) clustering with AaHD1 gene, suggesting it as a closest ortholog to *A. annua* HD-ZIP genes that is involved in the trichome development . Similarly, seven *M. citriodora* HD-ZIP-IV members (McHD-ZIP-IV-3, McHD-ZIP-IV-36, McHD-ZIP-IV-8, McHD-ZIP-IV-4, McHDZIP-IV-11, McHD-ZIP-IV-5, and McHD-ZIP-IV-2) clustered with AaHD8 gene. Thus, these close homologs of AaHD8 might be involved in the glandular trichome development in *M. citriodora*.

The HD-ZIV-IV transcription factors regulate the biosynthesis of metabolites such as cuticle wax, cutin, anthocyanin and flavanol biosynthesis **(Wang et al. 2020; Liu et al. 2025).** Therefore, apart from their role in the development of cuticular structure, these transcription factors may also directly regulate the biosynthesis secondary metabolites. Our co-expression analysis suggested that some of the HD-ZIP-IV transcription factor show close co-expression with ceratin terpene synthase genes family members. Additionally, few HD-ZIP IV genes coexpress with the DXS genes. These results suggested that HD-ZIP-IV family members might be involved in the regulation of terpenoid biosynthesis.

## Conclusions

The present study provided a more comprehensive and improved reference transcriptome assembly of M. citriodora, which will be useful for its functional genomics. The transcriptome-wide analysis led to the identification of HD-ZIP family transcription factors. A total of 41 putative McHD-ZIP IV sub-family transcription factors were identified. Based on the phylogenetic and expression analysis, 8 McHD-ZIP-IV were identified as putative homologs of known HD-ZIP TFs involved in the trichome development process in other species. Furthermore, based on gene-coexpression analysis with terpene synthase and DXS genes, several McHD-ZIP members were putatively predicted to be involved in terpenoid metabolism across different tissues, providing basis for studying the regulation of terpenoid metabolism through HD-ZIP TFs.

## Acknowledgments

Authors are thankful to CSIR, New Delhi, India for funding this activity under the project FBR020306. SA and KS acknowledge UGC for JRF and SRF. Director, CSIR-IIIM is acknowledged for providing necessary research facilities to carry out this work.

## Author’s contributions

Conceived and designed the experiments: PM; Performed the experiments: SA, KS, KP; Analyzed the data: SA, KS, AM, Contributed reagents/materials /analysis tools, PM, wrote the manuscript PM, SA, AM.

---

**S1:**
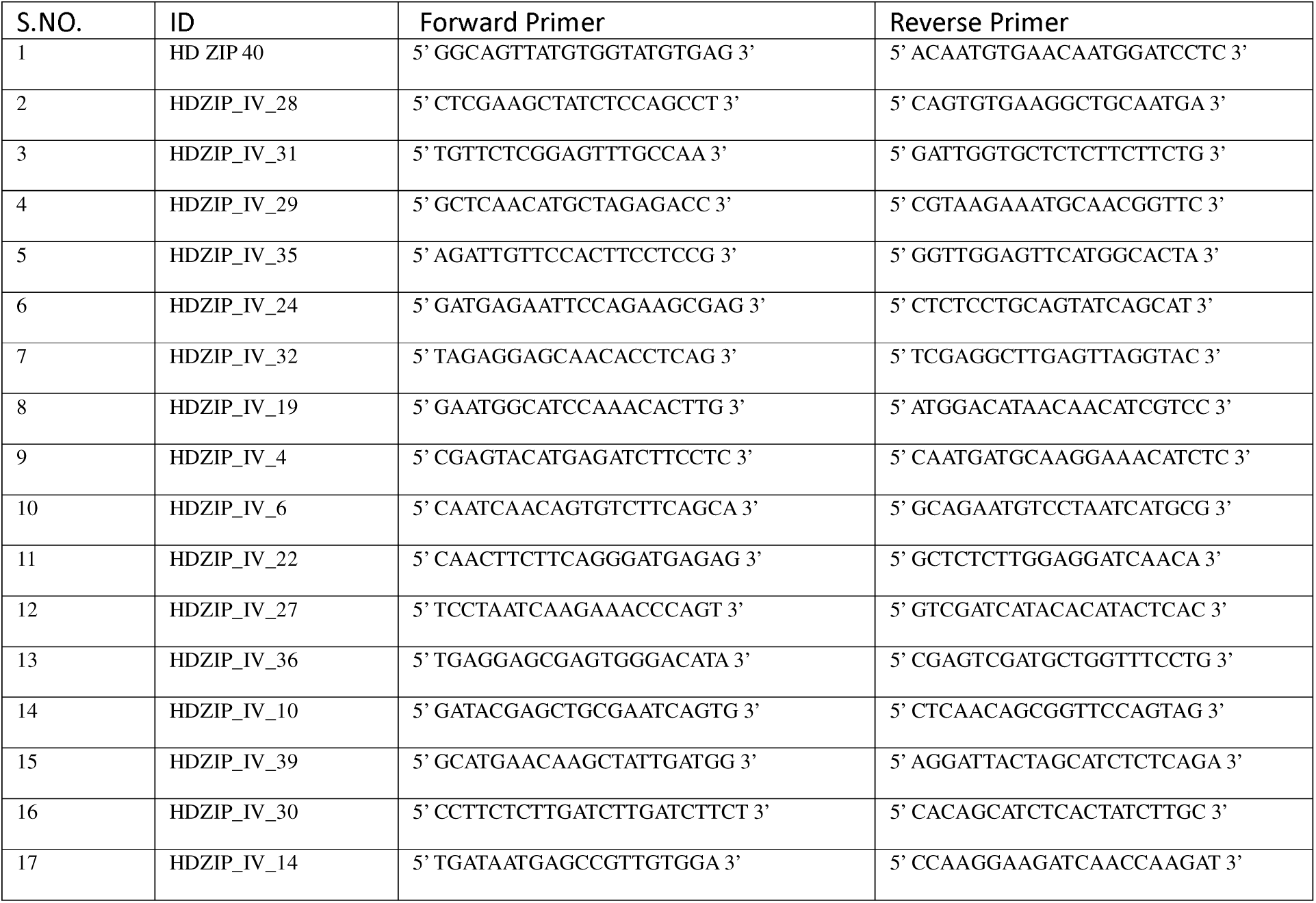
Supplementary table of primers used for RT-PCR analysis.

**S2:**
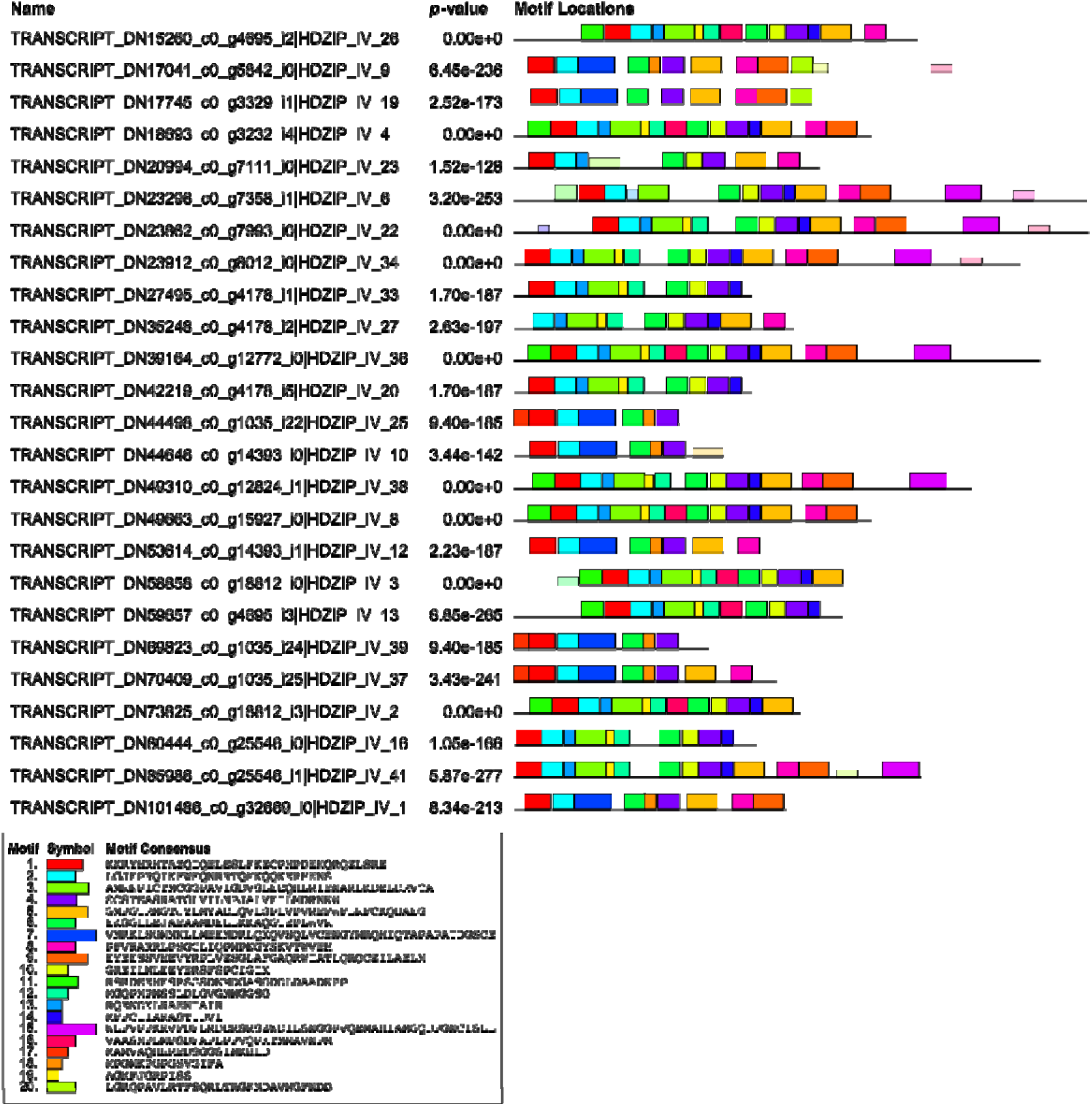
Graphical representation of motif distribution of the McHD-ZIP_IV members.

